# Microbial life under ice: metagenome diversity and *in situ* activity of Verrucomicrobia in seasonally ice-covered lakes

**DOI:** 10.1101/324970

**Authors:** Patricia Tran, Arthi Ramachandran, Ola Khawasik, Beatrix E. Beisner, Milla Rautio, Yannick Huot, David A. Walsh

## Abstract

Northern lakes are ice-covered for a large part of the year, yet our understanding of microbial diversity and activity during winter lags behind that of the ice-free period. In this study, we investigated under-ice diversity and metabolism of Verrucomicrobia in seasonally ice-covered lakes in temperate and boreal regions of Quebec, Canada using 16S rRNA sequencing, metagenomics and metatranscriptomics. Verrucomicrobia, particularly the V1, V3 and V4 subdivisions, were abundant during ice-covered periods. A diversity of Verrucomicrobia genomes were reconstructed from Quebec lake metagenomes. Several genomes were associated with the ice-covered period and were represented in winter metatranscriptomes, supporting the notion that Verrucomicrobia are metabolically active under ice. Verrucomicrobia transcriptome analysis revealed a range of metabolisms potentially occurring under ice, including carbohydrate degradation, glycolate utilization, scavenging of chlorophyll degradation products, and urea use. Genes for aerobic sulfur and hydrogen oxidation were expressed, suggesting chemolithotrophy may be an adaptation to conditions where labile carbon may be limited. The expression of genes for flagella biosynthesis and chemotaxis was detected, suggesting Verrucomicrobia may be actively sensing and responding to winter nutrient pulses, such as phytoplankton blooms. These results increase our understanding on the diversity and metabolic processes occurring under ice in northern lakes ecosystems.

## Originality and significance

Reduced ice cover on lakes is occurring worldwide, but there is only limited information on the biogeochemistry and microbiology under ice. This gap in knowledge limits our ability to understand and predict how changes in winter conditions will impact the ecology of lake ecosystems. In this study, we conducted the first meta-omics assessment of bacterial metabolism and gene expression patterns in seasonally ice-covered lakes. Previously uncharacterized lineages within the Verrucomicrobia were strongly associated with ice-covered conditions. Analysis of Verrucomicrobia genomes and gene expression patterns revealed a diversity of functional trait including the use of organic and inorganic energy sources and potential metabolic interactions with winter phytoplankton and zooplankton. The identification of winter-associated lineages and variable metabolic processes indicated that under-ice microbial communities may contribute uniquely to the ecology and nutrient cycling of seasonally ice-covered lakes. This study demonstrates the importance of studying the ice-covered period in the face of climate change and should spur future year-round investigations on microbial community structure and function in ice-covered freshwater ecosystems.

## Introduction

Many northern lakes are seasonally ice-covered for over 40% of the year (Weyhenmeyer *et al.*, 2011). During that time, these lakes are typically characterised by low light levels and primary productivity, leading to the traditional idea that they are “dormant” in winter (Bertilsson *et al.* 2013). However, recent studies describing phytoplankton and bacterial blooms as well as specialized microbial metabolism under ice have challenged this concept (Kankaala *et al.*, 2006; Twiss *et al.*, 2012; Bertilsson *et al.*, 2013; Bižić-Ionescu *et al.*, 2014; Beall *et al.*, 2016; Powers and Hampton, 2016). For example, light penetrating through ice and snow has been shown to fuel substantial primary production and blooms of low light and cold-adapted phytoplankton do occur under the ice (Twiss *et al.*, 2012; Üveges *et al.*, 2012). These phytoplankton blooms can fuel growth of other organisms, including heterotrophic bacteria (Bižić-Ionescu *et al.*, 2014). Moreover, unique niches can develop and persist in ice-covered lakes. Where ice-cover results in lower availability of labile organic substrates from phytoplankton or terrestrial input, volatile compounds such as methane and fermentation end-products produced from anoxic bottom waters or sediments can fuel microbial metabolism (Sundh *et al.*, 2005; Kankaala *et al.*, 2006). Chemolithoautotrophs may also be significant under ice since their sources of energy and carbon acquisition may not be as restricted by ice cover compared to phototrophs and heterotrophs (Auguet *et al.*, 2011). Despite these observations, our understanding of the structure and function of microbial communities under ice lags far behind that of the ice-free period (Bertilsson *et al.*, 2013; Powers and Hampton, 2016; Hampton *et al.*, 2017). To better understand the contribution of winter communities to lake metabolism and nutrient cycling, year-round investigation of the metabolic traits and activities of freshwater microorganisms is warranted.

Verrucomicrobia are ubiquitous in lakes, yet knowledge of their metabolism and ecology in freshwaters is limited compared to other bacterial groups (Newton *et al.*, 2011). The phylum is comprised of five orders, also referred to as subdivisions (V): Verrucomicrobiae (V1), Spartobacteria (V2), Pedosphaera (V3), Opitutae (V4), and Methylacidiphilum (V6) (Hedlund *et al.*, 1996; Sangwan *et al.*, 2004; Choo *et al.*, 2007; Hou *et al.*, 2008; Kant *et al.*, 2011). 16S rRNA gene surveys have identified all Verrucomicrobia subdivisions in lakes, however, only a few pelagic freshwater isolates exist, all belonging to V1 (Hedlund *et al.*, 1996). In general, while Verrucomicrobia are metabolically diverse, in aquatic ecosystems they are often associated with the degradation of carbohydrates (Martinez-Garcia *et al.*, 2012; Herlemann *et al*, 2013). For example, analyses of single cell-amplified genomes (SAGs) and metagenome-assembled genomes (MAGs) from coastal environments identified preferences for laminarin, xylan (Martinez-Garcia *et al.*, 2012), cellulose and chitin polymers (Herlemann *et al.*, 2013). Recently, comparative analysis of Verrucomicrobia MAGs from eutrophic Lake Mendota and dystrophic Trout Bog in Wisconsin, USA showed differences in the number and type of glycoside hydrolases (GHs) between the two systems, reflecting adaptations to local environments and carbon substrate availability (He *et al.*, 2017). Additionally, MAGs from freshwater reservoirs in Spain also suggested a preference for polysaccharides, but also identified genes for the light driven proton pump rhodopsin as well as genes for nitrogen fixation (Cabello-Yeves, Ghai, *et al.*, 2017). Perhaps most relevant from a life under ice perspective, a recent study identified abundant populations of Verrucomicrobia under the ice of Lake Baikal in Siberia, and these lineages also contained multiple polysaccharide degradation pathways in their genomes (Cabello-Yeves, Zemskaya, *et al.*, 2017). Given their widespread distribution across freshwater ecosystems and their prominence under ice, Verrucomicrobia may serve as a suitable model for investigating metabolic adaptations and lifestyle strategies associated with seasonally ice-covered lakes.

Here we present a study on Verrucomicrobia in seasonally ice-covered lakes in Quebec, Canada with a general aim of advancing our understanding of the genomic diversity and metabolic traits of bacterial communities residing under the ice of northern lakes. Using a combination of 16S rRNA gene sequencing, metagenomics, and metatranscriptomics we identified a wide diversity of Verrucomicrobia, including populations and MAGs strongly associated with ice-covered periods. The MAG-associated transcriptomes revealed a range of expressed metabolic genes under ice, including those for the use of phytoplankton and plant-derived organic compounds, chemolithotrophy, as well as motility and chemotaxis. Overall, our study supports the increasing recognition that the winter period represents a dynamic and metabolically important period for lakes despite ice cover.

## Results

### Verrucomicrobia diversity and distribution in Quebec Lakes

To investigate Verrucomicrobia abundance and diversity during ice-covered and ice-free periods of the year, we analyzed a 3-year time-series of bacterial 16S rRNA gene diversity from three seasonally ice-covered lakes located in the temperate (Lake Croche and Lake Montjoie) and boreal (Lake Simoncouche) regions of Quebec (**Fig. 1A**, **Table S1**). In total, we generated 16S rRNA data from 143 samples collected from the epilimnion and metalimnion, and corresponding to 6 winter time-points (January and February 2013, 2014, and 2015) and 8 summer time-points (June, July, August 2013 and 2014). The study lakes differed in several environmental characteristics, but were mainly differentiated by nutrient concentrations in a principal component analysis (**Fig. 1B**). The three lakes are distributed along a nutrient gradient, with Croche generally having lower total phosphorus (TP) than Montjoie and Simoncouche (**Fig. S1**, **Table S2**).

Verrucomicrobia were common across all lakes, but generally exhibited higher relative abundance of 16S rRNA gene sequences during ice covered period, particularly in Croche and Simoncouche (**Fig. 1C**). The average relative abundance of Verrucomicrobia was about 4-fold higher during ice cover (13%) compared to the ice-free period (4%) in Croche and about 2-fold higher in Simoncouche (10% compared to 5%). Although more abundant under the ice on average, Verrucomicrobia were highly dynamic in time with maximum observed values of 35.6 % and 27.1% in Croche and Simoncouche during winter time points. Verrucomicrobia in Croche were dominated by V1 and V3, while V4 was the major contributor in Simoncouche. In contrast to Croche and Simoncouche, Verrucomicrobia exhibited a similar mean relative abundance between ice-covered (4.8 %) and ice-free (6.2 %) periods in Montjoie. Maximum values in Montjoie were observed during summer time points and were comprised mostly of V6. Although relative abundance differed between lakes, these results demonstrate Verrucomicrobia are often associated with ice-covered conditions, making them an intriguing group with which to investigate genomic and metabolic adaptations to life under ice.

**Fig. 1.**
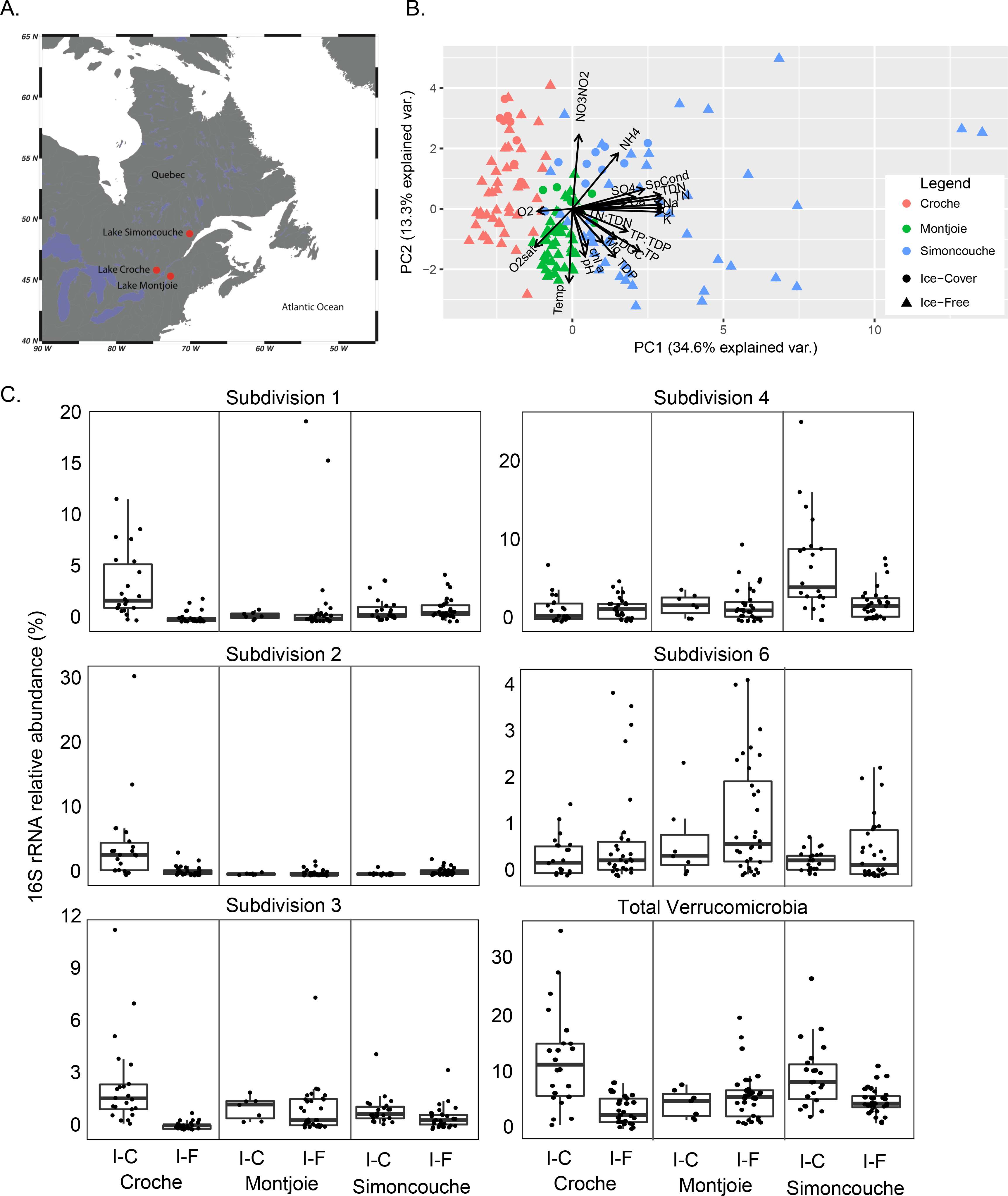
**A.** Locations of Lake Croche, Lake Montjoie and Lake Simoncouche in temperate and boreal regions of Quebec, Canada. **B.** Environmental variability of lake samples based on principal component analysis of 21 physiochemical variables. **C**. Relative abundance of Verrucomicrobia in the 16S rRNA amplicon datasets organized by lake and season. I-C and I-F refer to ice-covered and ice-free periods, respectively. Mid-lines represent median, upper and lower boundaries are 25% quartiles, and whiskers represents scores outside the middle 50%.

### Verrucomicrobia MAG diversity and gene transcription

Verrucomicrobia genome diversity was investigated in a metagenome co-assembly of 24 samples collected over the 3-year times-series from the three lakes. Following metagenomic binning, we identified 54 Verrucomicrobia MAGs representing all 5 subdivisions (V1 to V4 and V6) previously observed in the 16S rRNA gene survey (**Fig. 2A**). To investigate the distribution of MAGs across lakes and seasons, we performed a Canonical Correspondence Analysis (CCA) of MAG coverage across samples, constrained using 14 environmental variables (see methods for definitions of abbreviations). In the resulting CCA, samples were differentiated along two primary axes (**Fig. 2B**). Croche samples were separated from Montjoie and Simoncouche samples along CCA axis 1, which correlated with nutrient concentrations (*i.e.* TP and TN). Ice-covered samples were generally separated from ice-free samples along CCA axis 2, which differentiated samples from the ice-covered and ice-free period, with the exception of a single summer Montjoie sample, which clustered with winter samples. A clear separation of MAGs along both axes of the ordination was observed, revealing lake and seasonal preferences (**Fig. 2C**). Interestingly, MAGs within a subdivision, and even those closely related within a subdivision, did not cluster together in the CCA, suggesting niche diversity among members of the same subdivision; taxonomy clearly does not reflect niche preference.

**Fig. 2.**
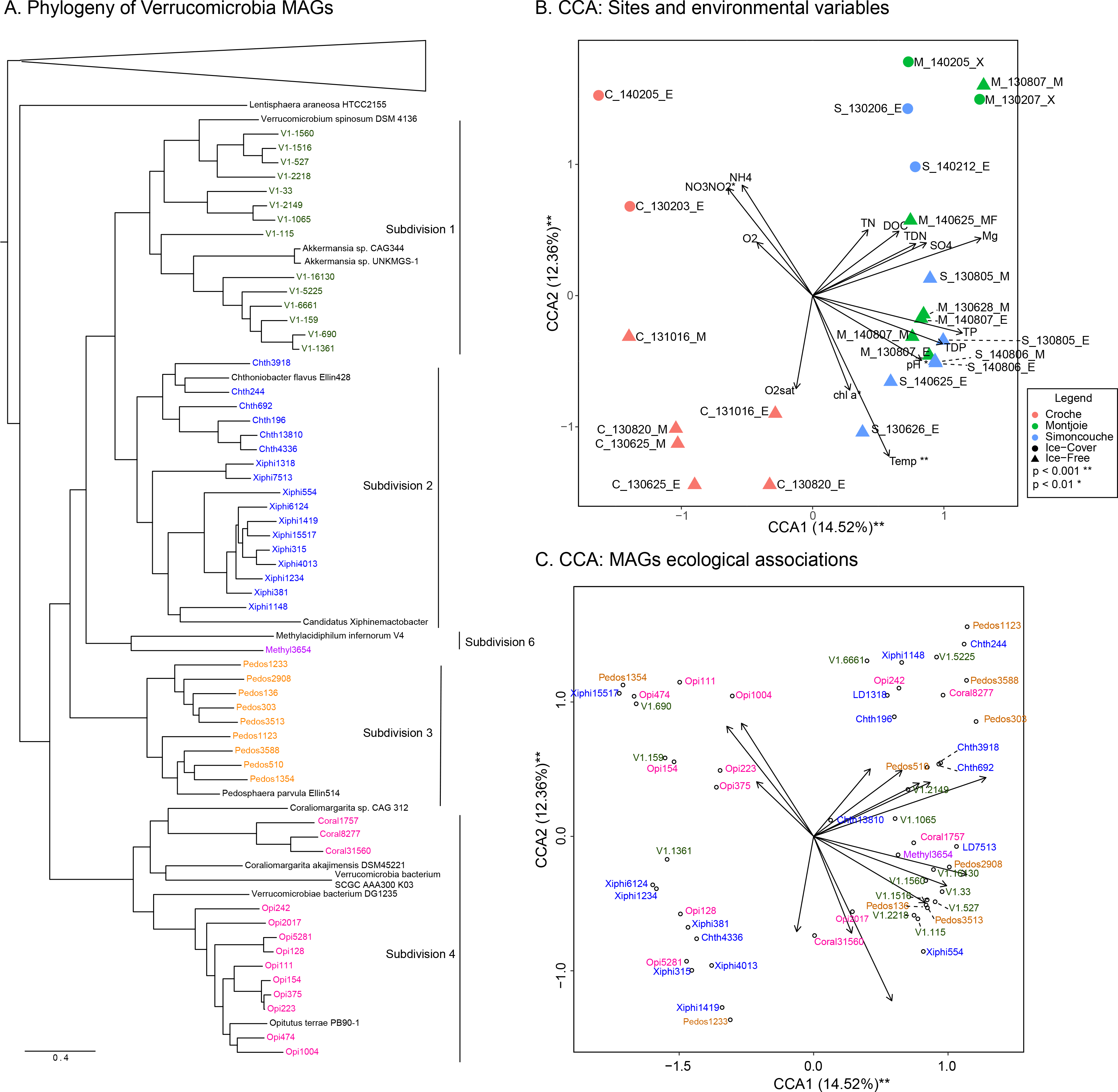
**A.** Concatenated protein phylogeny of sampled lake MAGs and Verrucomicrobia reference genomes. **B. and C.** CCA of the 54 MAGs constrained to 14 environmental variables (arrows). The 24 samples are shown in the ordination space in panel b, whereas the 54 MAGs are shown in panel c. Abbreviations for environmental variables are described in Table S2.

Fifteen MAGs of high completeness and low contamination were selected for further analyses (**Table 1**). These MAGs represented four subdivisions (V1 to V4). The MAGs were distributed across the CCA plot, and are therefore representative of the observed phylogenetic diversity, biogeography, and seasonal associations of Verrucomicrobia MAGs. A concatenated protein phylogeny that included 43 additional MAGs from Wisconsin lakes (He *et al.*, 2017), Lake Baikal (Cabello-Yeves, Zemskaya, *et al.*, 2017) and the Tous and Amadorio reservoirs in Spain (Cabello-Yeves, Ghai, *et al.*, 2017) was generated in order to place the Quebec MAGs in the context of known Verrucomicrobia genomic diversity (**Fig. 3A**). The phylogeny presented in Figure 3A was inferred from a concatenated alignment of four of the five proteins used in He et al., 2017, but exhibited a similar topology to a phylogeny generated based on hundreds of proteins (**Fig. S2**). MAGs were too distantly related to calculate average nucleotide identity (ANI), but average amino acid identity between Quebec MAGs and those from other locations ranged between 91-49% (**Table S3**). For the most part, Quebec MAGs exhibited <70% AAI with other MAGs and therefore represent new lineages for which genome sequence data is now available. Although common across Quebec lakes, fragment recruitment of metagenomes originating from Wisconsin lakes, Lake Baikal, and the Spanish reservoirs demonstrated that the Verrucomicrobia populations represented by the MAGs are relatively rare in these other freshwater systems (**Table S4**).

**Table 1.** Genome characteristics of the 15 Verrucomicrobia MAGs, recovered from Quebec lakes metagenomes co-assembly.

| Subdiv. | MAG | Sequence<br>length<br>(Mbp)<sup>a</sup> | N50<sup>a</sup> | Coverage<sup>b</sup> | GC<br>Content<br>(%)<sup>a</sup> | Gene<br>Count<sup>a</sup> | Complete-<br>ness (%)<sup>a</sup> | Contami-<br>nation<br>(%)<sup>a</sup> | Estimated<br>bin size<br>(Mbp)<sup>c</sup> |
| --- | --- | --- | --- | --- | --- | --- | --- | --- | --- |
| 1 | V1-33 | 4.08 | 49467 | 11.9 | 63.31 | 3225 | 78.24 | 0.68 | 5.18 |
| 1 | V1-115 | 4.14 | 40207 | 9.9 | 61.77 | 3420 | 82.07 | 4.55 | 4.81 |
| 1 | V1-159 | 2.63 | 31999 | 16.3 | 62.55 | 2373 | 95.92 | 0 | 2.74 |
| 1 | V1-690 | 2.24 | 17084 | 12.3 | 60.46 | 2015 | 70.61 | 2.86 | 3.08 |
| 1 | V1-1361 | 2.25 | 11724 | 23.8 | 60.29 | 2055 | 72.93 | 0.68 | 3.06 |
| 2 | Chth-244 | 3.24 | 18974 | 13 | 65.1 | 3122 | 83.28 | 2.2 | 3.80 |
| 2 | Chth-196 | 2.79 | 35078 | 15.1 | 58.27 | 2263 | 70.16 | 0 | 3.98 |
| 2 | Xiphi-554 | 1.66 | 24984 | 10.4 | 42.77 | 1576 | 95.95 | 0.71 | 1.72 |
| 2 | Xiphi-315 | 1.42 | 22457 | 38.9 | 54.51 | 1400 | 87.73 | 2.03 | 1.59 |
| 3 | Pedos-303 | 4.47 | 17418 | 9 | 67.77 | 3693 | 89.7 | 2.7 | 4.85 |
| 3 | Pedos-1123 | 2.73 | 11423 | 12.6 | 52.1 | 2471 | 95.05 | 2.59 | 2.80 |
| 3 | Pedos-510 | 2.51 | 16489 | 25.8 | 54.52 | 2380 | 82.43 | 0.34 | 3.03 |
| 4 | Opi-242 | 1.19 | 34025 | 63.8 | 54.32 | 1116 | 57.77 | 0 | 2.06 |
| 4 | Opi-128 | 1.85 | 18982 | 107.1 | 42.47 | 1684 | 77.74 | 0.68 | 2.36 |
| 4 | Opi-474 | 2.81 | 11480 | 51.6 | 65.6 | 2690 | 92.17 | 4.11 | 2.92 |

<sup>a</sup> Calculated using CheckM
<sup>b</sup> Calculated using MetaWatt
<sup>c</sup> Calculated as (sequence length/completeness)\*(100-contamination)

As revealed by their wide distribution in the CCA ordination, the MAGs exhibited complex distribution patterns across Quebec lakes (**Fig. 3B**). To investigate gene expression patterns of the Verrucomicrobia MAGs during ice-covered and ice-free periods in Quebec, a temporally overlapping metatranscriptomic times-series was mapped to the MAGs, providing a view of their transcriptional activity across lakes and seasons. A significant number of transcripts were observed for all MAGs, and for the most part reflected MAG distributions across lakes and seasons (**Fig. 3B**). In the following section, we analyzed the distribution, metabolic gene content and gene expression patterns of Verrucomicrobia MAGs with the objective of providing insights into metabolic diversity and activity associated with ice-covered conditions in northern lakes.

**Fig. 3.**
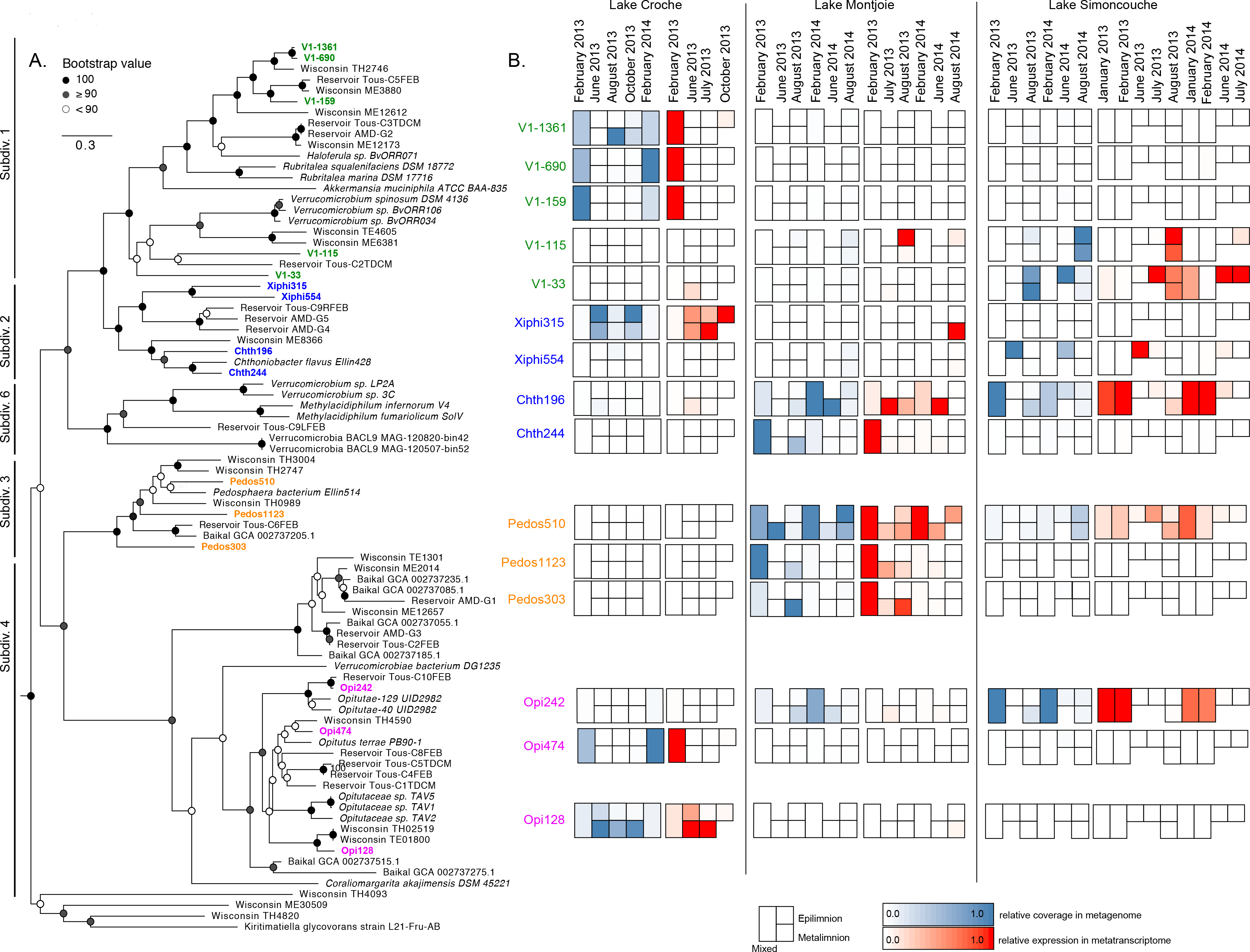
**A.** Concatenated maximum likelihood phylogeny, using 100 boostraps, of selected Verrucomicrobia MAGs and reference genomes. **B.** Heatmap of the relative MAG abundance based on coverage (blue) and relative gene expression based on the number of transcripts recruited (red) to each MAG. Rectangular boxes represent ice-covered samples, square boxes represent epilimnetic (top squares) and the metalimnetic (bottom squares) samples.

### Ice cover associated MAGs within V1

A striking association with winter conditions was observed within certain V1 MAGs. Particularly, three MAGs (V1-159, V1-690, and V1-1361) were associated with ice-covered conditions in Croche. These MAGs formed a clade with other freshwater MAGs but were quite unique, sharing between 51-72 % AAI (**Fig 3A**, **Table S3**). V1-690and V1-159 were exclusive to winter periods both at the level of the genome and the transcriptome (**Fig. 3B**), while V1-1361 was also identified in the summer. A total of 3,983 orthologs were identified among these three MAGs and 868 were common to all (**Fig. 4A**). Similar to other Verrucomicrobia, the winter-associated V1-159 and V1-690 MAGs contained numerous glycoside hydrolase (GH) genes (**Table S5**). Expression of 33 of 34 GH genes was detected in V1-159. The GHs of highest expression were GH29 (alpha-L-fucosidase) and GH16 (substrate specificity undetermined). In contrast, expression was only detected for 12 of 54 GH genes in V1-690, suggesting other modes of carbon and energy metabolism may be important for this population. Indeed, four proteins annotated as monooxygenases were specific to V1-690 and were relatively highly expressed. Two were annotated as limonene 1,2-monooxygenase and a third as 1 alkane 1-monooxygenase/p-cymene monooxygenase (**Fig. 4B**). Limonene and cymene are plant-derived aromatic compounds, suggesting that these proteins are involved in accessing terrestrial organic matter. In addition, we identified a genomic region encoding the Sox system (*soxXYZABC*) together with two c-type cytochromes (**Fig. 4C**). The Sox system is associated with the use of the reduced sulfur compound thiosulfate as an electron donor in energy metabolism (Ghosh and Dam, 2009). All subunits were expressed during winter except for *soxZ*, suggesting the potential for lithotrophic sulfur oxidation during ice-covered periods.

**Fig. 4.**
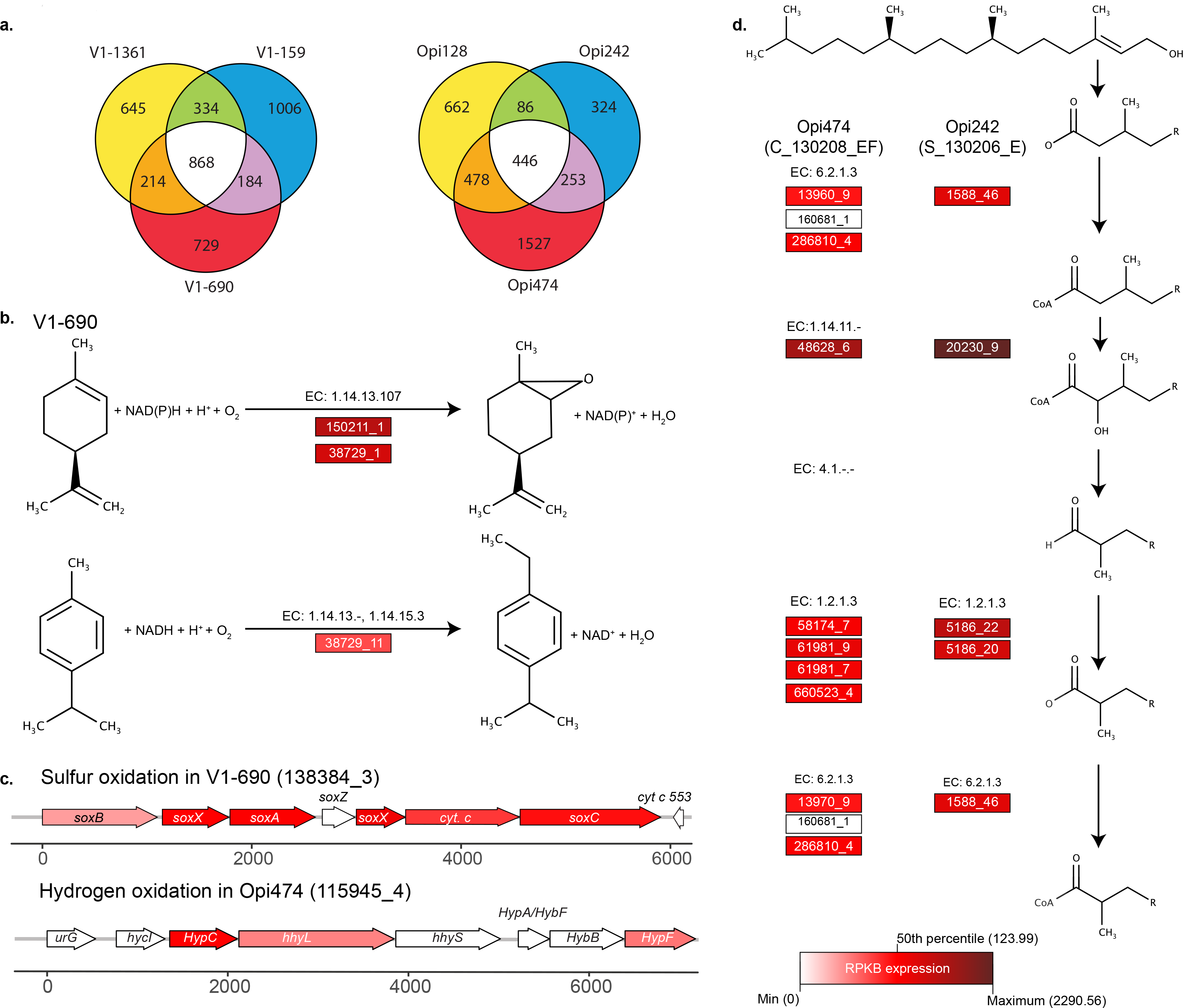
**A.** Venn diagrams of ortholog distributions between MAGs in V1 (left panel) and V4 (right panel). **B.** Enzymatic reaction for limonene (top) and cymene (bottom) degradation identified in V1-690. **C.** Gene clusters involved in sulfur oxidation in V1-690 and hydrogen oxidation in Opi-474. **D.** The α-oxidation pathway for phytol degradation in Opi-474 and Opi-242. For panels **b** and **e**, E.C. numbers are shown, and shaded boxes represent RPKB during winter. Numbers inside the boxes represent “ScaffoldNumber_Loci”.

### Ice cover associated MAGs within V4

Specific winter association was observed for V4 MAGs. Two MAGs (Opi-128 and Opi-474) were phylogenetically related to MAGs from Trout Bog and the Tous reservoir. Opi-128 exhibited 81 % AAI with TH02519 and TH01800 and Opi-474 exhibited 82 % AAI with TH4593, while Opi-128 exhibited 91 % AAI with Tous-C10FEB (**Fig. 3A**, **Table S3**). Opi-474 and Opi-242 exhibited winter association in Croche and Simoncouche, respectively (**Fig. 3B**). Opi-128 was more commonly associated with summer. A total of 3,776 orthologs were identified among the three genomes and only 11% were shared among all MAGs (**Fig. 4A**), indicating substantial genomic diversity among V4 MAGs. Winter associated Opi-474 and Opi-242 contained 70 and 19 GH genes, respectively (**Table S4**). Compared to V1 winter MAGs, the abundance of GH transcripts was elevated in the Opi-474 transcriptome. Among the expressed GHs were GH78 genes involved in the use of rhamnose-containing polysaccharides.

A common feature of Opi-474 and Opi-242 was the expression of a predicted fatty acid α-oxidation II pathway (**Fig. 4D**). Fatty acid α-oxidation is implicated in the metabolism of phytol, a long chain alcohol constituent of chlorophyll (Jansen and Wanders, 2006). Phytol is first converted to phytanoyl-CoA, which then enters the α-oxidation pathway where a methyl group at the C-3 position is removed before passage to the more common β-oxidation pathway (Jansen and Wanders, 2006). Both genomes encoded proteins of the phytanoyl-coA dioxygenase family, which hydroxylates themethyl-branched fatty acid preparing it for downstream cleavage and passage to the β-oxidation pathway. A putative pathway containing alcohol and aldehyde dehydrogenases necessary for transforming phytol to phytanoyl-CoA was also identified. The presence of a putative α-oxidation pathway and upstream steps for introducing phytol may allow these organisms to scavenge carbon and energy from chlorophyll degradation products.

Evidence for chemolithotrophic energy conservation was observed in V4. A complete gene cluster associated with aerobic hydrogen oxidation was specifically identified in Opi-474, including genes encoding a Group 1d oxygen-tolerant hydrogenase (Greening *et al.*, 2016) and associated maturation and nickel incorporation proteins (**Fig. 4C**). Expression of a number of these genes was detected under the ice in Croche, suggesting the use of hydrogen as an electron donor in energy metabolism.

### Complex seasonal dynamics in V2 and V3 MAGs

Compared to V1/V4 MAGs, V2 and V3 MAGs exhibited more complex distribution patterns across lakes and seasons. In V3, Pedo-303 and Pedo-1123 branched deeply within V3, and were present throughout the year in Montjoie, while Pedo-510 formed a clade with MAG recovered from Trout Bog and was present throughout the year in Montjoie and Simoncouche (**Fig. 3**). V3 Quebec MAGs exhibited less than 66 % AAI with MAGs from Trout Bog and the Tous reservoir.

The most complex seasonal patterns were observed for the four V2 MAGs. Two MAGs (Xiphi-315 and Xiphi-554) were members of the *Xiphinematobacter*. Xiphi-315 was common but variable in Croche summer samples, while Xiphi-554 was restricted tothe June epilimnion in Simoncouche. Quebec *Xiphinematobacter* MAGs (Xiphi-315 and Xiphi-554) were within the same clade as three MAGs from the Tous and Amadorio reservoirs, but shared at most 63% AAI with them, reflecting high genome variation between MAGs from Quebec lakes and other freshwater environments. Interestingly, Xiphi-554 was the most widely distributed in other freshwater systems based on metagenome fragment recruitment patterns (**Table S4**)

The two other V2 MAGs (Chth-244 and Chth-196) were members of the *Chthoniobacter*. Chth-196 was the most broadly distributed MAG of all in Quebec lakes, being common in Montjoie and Simoncouche but exhibiting high gene expression levels in all. However, Chth196 was the only MAG that did not recruit any other freshwater metagenome reads (**Table S5**), indicating very unique populations in Quebec lakes. Chth-244 was common throughout the year in Montjoie although transcripts were only detected during ice cover.

With respect to physiological adaptations to life under ice, Pedo-303 (V3) and Chth-196 (V2) are of interest because differences in the number of transcripts between summer and winter was detected for both MAGs, providing insight into metabolic responses to different seasons. Although Pedo-303 and Chth-196 were distantly related from a phylogenetic perspective, they exhibited some similarities at the metabolic level. For example, ammonia and urea appear to be important nitrogen sources for both groups. Urea transport proteins and urease subunits were identified in Chth-196 and generally had higher expression during winter than summer in Simoncouche. Urea transport and urease genes were also present in Pedo-303 and more transcripts were detected during ice cover than the ice-free period in Montjoie (**Fig. 5C**). Furthermore, Pedo-303 had genes implicated in ammonia transport and assimilation, including three copies of the nitrogen regulatory proteins P-II1, two Amt family ammonium transporters, and two glutamine synthetases, which had higher levels of expression in the winter (**Fig. 5D**). In Chth-196, three Amt family transporters, one glutamine synthetase, and three nitrogen regulatory proteins P-II-1 were found, and these genes were expressed more in Simoncouche during ice cover, but more in Montjoie during the summer.

Pedo-303 and Chth-196 both encoded genes for the use of a range of organic carbon compounds. In Pedo-303, rhamnose and xylose utilization genes were expressed at both time points, but transcript abundance was higher in the winter. In Chth-196, genes for rhamnose and ribose transport and degradation were identified and expressed during winter. The use of glycolate, a photorespiration product of phytoplankton, was also suggested by the presence of a glycolate utilization operon (Fe-S subunit, FAD-binding subunit, oxidase) in Pedo-303. Transcripts for glycolate use were only detected during winter in Montjoie.

Another notable observation within V2 and V3 MAGs was the relatively high number of transcripts for genes involved in motility and chemotaxis. Genes for flagellar biosynthesis were identified in numerous MAGs (Chth-196, Chth-244, Opi-474, Pedo-510 and Xiphi-554), but evidence for active use of the flagellar machinery was found in Chth-196 and Pedo-510 only. In Chth-196, transcripts encoding flagellar motility proteins were generally more abundant in summer in Montjoie, but in winter in Simoncouche (**Fig. 5A**). In addition, Chth-196 encoded genes involved in chemotaxis behaviour were more abundant in winter than summer in Simoncouche (**Fig. 5B**).

**Fig. 5.**
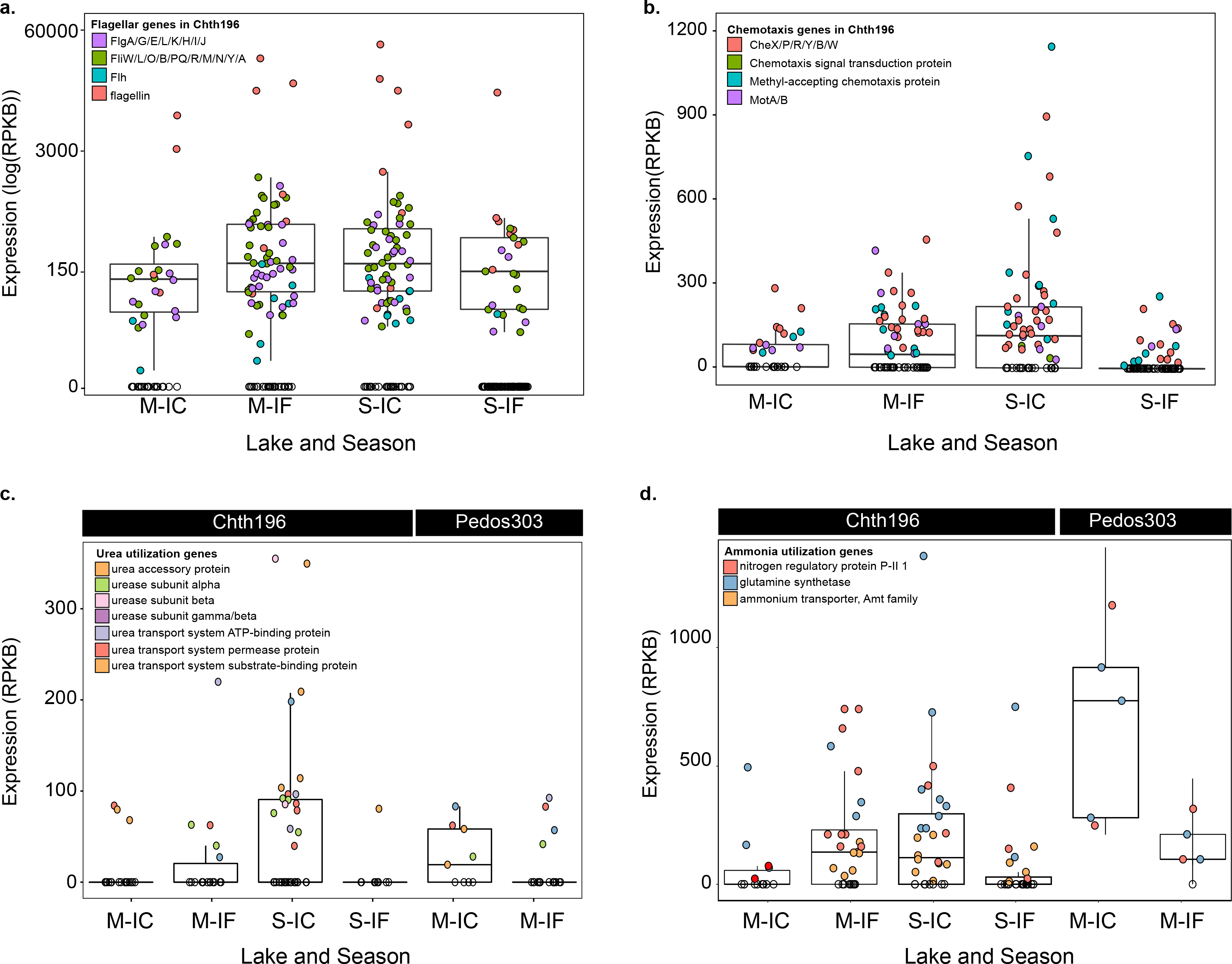
**A.** Expression of Chth-196 genes involved in flagellar motility. **B.** Expression of Chth-196 genes involved in chemotaxis **C.** Expression of genes in the urease gene cluster in Chth-196 and Pedo-303 **D.** Genes involved in ammonia utilization and degradation in Chth-196 and Pedo-303. Since Chth-196 is present in Montjoie and Simoncouche, but Pedos-303 is only present in Montjoie, only those lakes are shown. Genes for which expression was positive (Reads per kilo bases per billions reads ‑ RPKB) are highlighted with colour.

## Discussion

In this study, we conducted the first meta-omics assessment of bacterial metabolism and gene expression patterns in seasonally ice-covered lakes. In combination with other recent metagenomic studies (Cabello-Yeves, Ghai, *et al.*, 2017; Cabello-Yeves, Zemskaya, *et al.*, 2017; He *et al.*, 2017), our findings contribute to the emerging view that Verrucomicrobia is an important, but previously overlooked component of lake microbial ecosystems. The association of Verrucomicrobia MAGs with ice-covered conditions suggests certain populations exhibit a winter preference. However, an alternative explanation for greater under-ice abundance is that Verrucomicrobia prefer as habitat the deeper hypolimnetic regions of northern lakes, and thus they could appear winter-associated following autumn mixing of the water column. Although we cannot rule out this possibility, it seems unlikely given that sampling occurred several months following the onset of fall overturn. Furthermore, we did not observe evidence for hypolimnetic preferences via anaerobic metabolism in the Verrucomicrobia MAGs despite the fact that Croche and Simoncouche have anoxic hypolimnia during summer. However, Verrucomicrobia have been observed in the oxic hypolimnion of the Tous and Amadorio reservoirs (Cabello-Yeves, Ghai, *et al.*, 2017). Moreover, studies of the sub-ice microbial communities of deep Lake Baikal have shown Verrucomicrobia are among the most dominant groups at the surface (Cabello-Yeves, Zemskaya, *et al.*, 2017) and that their abundances are linked to diatom blooms (Bashenkhaeva *et al.*, 2015). Hence, it appears that Verrucomicrobia populations represented by our MAGs are resident members of the microbial community in ice-covered periods in northern lakes, and knowledge of their genomic and metabolic traits can contribute to our understanding of lake metabolism and nutrient cycling in winter.

Ice-covered conditions may favour microbes that have low resource requirements (*i.e.* oligotrophs) and small genome size is a common adaptation to oligotrophic conditions (Carini *et al.*, 2012; Neuenschwander *et al.*, 2017). Previous analysis of Verrucomicrobia MAGs have shown a wide range in genome sizes (Cabello-Yeves, Ghai, *et al.*, 2017). We speculated that winter-associated MAGs may be specialized to low carbon/energy conditions and that this would manifest in overall smaller genomes compared to those associated with the ice-free period. However, we did not observe generalizable differences in estimated genome size across seasons within Quebec lakes. Rather, we observed that overall estimated genome size was on average smaller in Verrucomicrobia MAGs from more oligotrophic lakes (Quebec, Lake Baikal, and freshwater reservoirs) compared to eutrophic lakes (Mendota and Trout Bog). Wisconsin MAGs had the largest average estimated genome size in Subdivision 1, 2 and 3 compared to Quebec, Baikal, Tous and Amadorio MAGs. The largest average estimated genome size in V4 was in Tous and Amadorio MAGs, while the smallest was among Quebec MAGs. Based on our results, small genome size does not seem to be distinguishing characteristic of winter association, at least for Verrucomicrobia, but instead reflects adaptations to oligotrophic freshwater environments in general.

### Verrucomicrobia-phytoplankton coupling

Phytoplankton can persist and form transient blooms during ice cover (Twiss *et al.*, 2012) and a number of studies have implicated Verrucomicrobia in the degradation of phytoplankton-derived carbohydrates(Paver and Kent, 2010; Parveen *et al*, 2013), including in ice-covered lakes (Bižić-Ionescu *et al.*, 2014; Beall *et al.*, 2016). For example, Verrucomicrobia were among the main bacteria associated with diatom-dominated under ice blooms in Lake Baikal (Bashenkhaeva *et al.*, 2015). Furthermore, Verrucomicrobia OTUs, including *XipA1/XipB1* (V2) and *Opitutacea* (V4) were strongly correlated with algal carbon in Finnish subarctic lakes (Roiha *et al.*, 2016). Finally, Verrucomicrobia (*Verrucomicrobiae* (V1) and *Opitutacea* (V4)) abundance significantly increased following the release of extracellular polymeric substances released by diatoms in intertidal zones (Bohórquez *et al.*, 2017)

Numerous traits of the Quebec lake Verrucomicrobia suggest a capacity to couple growth to phytoplankton, including winter phytoplankton blooms. Based on 16S rRNA gene analysis, Verrucomicrobia subdivisions were commonly abundant in both the free-living and particle-attached fractions of the community, indicating an ability to switch between these lifestyles. During winter, cells may persist in a free-living state but also associate with phytoplankton during bloom onset and progression. If so, then the MAGs analysed here originated from the free-living persisters. Interestingly, GH genes were identified in all MAGs, with no strong correlation between estimated genome size and number of GH found. However, we found a wider diversity of GHs was expressed in winter compared to summer, perhaps in readiness for persister cells to quickly respond to carbohydrate that becomes available, both from phytoplankton or terrestrial sources. Expression of genes for the use of phytoplankton-derived saccharides including fucose and rhamnose were identified, suggesting that Verrucomicrobia interact with phytoplankton during winter. Future studies comparing gene expression patterns between particle-attached and free-living cells, would be informative in understanding the metabolic shifts that occur during these transient bloom events.

Intriguingly, expression of a glycolate degradation operon was detected in ice cover associated MAGs from Lake Montjoie. Glycolate is a photorespiration product (Fogg, 1983) that is typically produced by phytoplankton growing under high light stress (Parker and Armbrust, 2005; Davis *et al.*, 2013). Glycolate-utilizing bacteria have been shown to be transcriptionally responsive to phytoplankton blooms (Lau *et al.*, 2007). Glycolate production during winter may be unexpected owing to low light penetration through snow and the absence of light stress (Maxwell *et al.*, 1994). However, low-light adapted phytoplankton may have a lower tolerance to light overall. Rapid increases in light intensity due to blowing/melting snow could lead to light stress, and a corresponding pulse of glycolate into the environment. Another possibility is that glycolate produced during the ice-free period is relatively long-lived in the water column. Glycolate is present in the ocean year-round, and can account for as much as 33% of the dissolved organic carbon pool (Carlson and Ducklow, 1996; Leboulanger *et al.*, 1997). In any case, it is believed that the main role of glycolate in heterotrophic metabolism is as an energy source (Wright and Shah, 1977; Edenborn and Litchfield, 1987). Recently, a study of carbon utilization in temperate lakes showed that a relatively larger portion of phytoplankton-derived carbon was allocated to respiration (and hence energy conservation), while terrestrial carbon was allocated to biosynthesis (Guillemette *et al.*, 2016). Therefore, glycolate may be an important contributor to the maintenance energy of a persisting cell or could facilitate growth by serving as an energy source, fuelling the subsequent assimilation of terrestrial carbon in winter.

Phytoplankton blooms and resource availability under ice are often patchy. Motility and chemotaxis may be useful adaptations to efficiently exploit hotspots of organic and inorganic nutrients (Stocker and Seymour, 2012). This idea is supported by a number of Verrucomicrobia MAGs that expressed genes for flagella and chemotactic abilities. Of particular interest were two closely related winter MAGs from Montjoie (Chth-244) and Simoncouche (Chth-196). Flagella and chemotaxis gene expression was evident in the Montjoie MAG. Despite the presence of flagella and chemotaxis genes in the Simoncouche MAG, only the flagella genes were detectably expressed. The difference in chemotactic activity may reflect the difference in lake physical conditions; while Montjoie is unstratified in winter, Simoncouche is inversely stratified. Hence, the ability to detect and respond to chemical gradients may be more advantageous in stratified systems. Operating at the micro-scale rather than meters, this notion of motility towards hotspots of nutrients is analogous to findings in phytoplankton motility. For example, in a 20-year study of phytoplankton functional traits in an ice-covered lake in Germany, authors found that during mild winters a mixed water column favoured non-motile phytoplankton (Özkundakci *et al.*, 2016), as in Montjoie. Therefore, the difference in water column mixing between lakes is potentially reflected in bacterial functional traits as well.

### Scavenging of the phytol moiety of chlorophyll

In addition to responding to algal growth, bacterial scavenging of dead and degraded phytoplankton and other detritus may be an important metabolic strategy for life under ice. Labile algal carbohydrates and proteins would likely be the first biomolecules scavenged from the environment, but carbon-rich lipids may also sustain growth during winter. In freshwater reservoirs, Verrucomicrobia metagenomes contained several enzymes (aryl sulfatases, beta-galactosidases and sialidases) for degradation of glycosphingolipids and are suggested to be involved in the degradation of plant or algae derived sulfur-containing lipids (Cabello-Yeves, Ghai, *et al.*, 2017). In our study, winter MAGs encoded for a predicted α-oxidation pathway. The α-oxidation pathway is implicated in the degradation of phytol, the long-chained alcohol moiety of chlorophyll. Early studies showed growth of bacteria on phytol as a sole source of carbon and energy (Hoag *et al.*, 1969), however the ecological significance of this metabolism is unknown. The concentration of free phytol in the water column may be low, but we can envision microenvironments where phytol concentrations are elevated. One micro-environment may be formed by the fecal pellets of zooplankton. Early studies have found that the fecal pellets of zooplankton, specifically coccoliths feeding on an algal-rich diet, were rich in phytanic acid, the end product of phytol oxidation that feeds into the α-oxidation pathway (Moussa, 1988). The phytanic acid-rich particles would be available in the water column for consumption by bacteria possessing the α-oxidation pathway

Additionally, it is possible that phytol may accumulate in particles that sink into the anoxic hypolimnion of lakes. Since the α-oxidation pathway requires oxygen (through the essential phytanoyl CoA hydroxylase), phytol may accumulate under anoxic conditions, although anaerobic degradation of phytol in sediments has been reported (Rontani *et al.*, 1999). Nevertheless, during breakdown of stratification in autumn, phytol accumulated in the hypolimnion could serve as a carbon reserve supporting microbial metabolism throughout the winter. A recent study in Lake Simoncouche has shown that zooplankton are able to store fatty acids from phytoplankton to survive over the winter (Grosbois *et al.*, 2017). Similarly, bacteria might be able to use zooplankton and phytoplankton-derived lipids (e.g. phytanic acid) to maintain baseline metabolism under ice. These observations point to the complex relationships among bacteria, phytoplankton, and zooplankton occurring under the ice in freshwater lakes.

### Chemolithotrophy under ice

Chemolithotrophic growth is traditionally thought to be restricted to sediments or the chemocline of stratified lakes where there is a sufficient supply of reduced inorganic compounds for metabolism. Here we detected the expression of sulfur and hydrogen gene clusters, suggesting that chemolithotrophic energy metabolism may be employed by bacteria under the ice. Similar sulfur oxidation genes were also recently reported in *Beta-proteobacteria* MAGs from Lake Baikal (Cabello-Yeves, Zemskaya, *et al.*, 2017). Heterotrophic organisms capable of supplementing their energy demand using inorganic compounds would have an advantage over those that cannot if the availability of organic carbon is limited in winter. A wide diversity of so-called “heterotrophic sulfur-oxidizing” bacteria are found in marine systems (Teske *et al.*, 2000). In these cases, sulfur oxidation allows bacteria to allocate organic carbon for biosynthesis instead of respiration, allowing them to thrive in a wider range of habitats (Teske *et al.*, 2000; Podgorsek *et al.*, 2004). Similarly, the winter water column availability of H_2_ (previously restricted to the anaerobic hypolimnion), combined with the availability of oxygen under ice, may allow aerobic hydrogen-oxidizing to use this energy rich fermentation end-product for growth or maintenance energy.

### Urea as an important N source in lake

The observation of urea utilization genes in our MAGs further suggests the potential for trophic interactions. Although urea is usually present in lakes at ambient concentrations below 1 μM-N, it can contribute 50% or more of the total N used by planktonic communities (Solomon et al., 2010). Interestingly, urea availability is predicted to be regulated mainly by the decomposition of algae under anoxic conditions (e.g. hypolimnion/sediments), followed by redistribution in the water column (Bogard *et al.*, 2012) and thus potentially serving as a valuable nitrogen source in winter. All urease subunits were found and expressed in the winter in Chth-196 and Pedo-303. A recognized source of N for polar phytoplankton, the importance of urea for bacteria has received less attention. In the arctic, urea utilization was detectable in the prokaryotic size fraction, but not that corresponding to the phytoplankton fraction, showing the potential importance of urea for cold-adapted metabolism, and adaptation to low energy environments (Alonso-Sáez *et al.*, 2012). In Lake Baikal, urea utilization genes were found in *Cyanobacteria, Acidobacteria, Nitrospearea, Beta-proteobacteria and Thaumarchaeota* MAG (Cabello-Yeves, Zemskaya, *et al.*, 2017). The expression of the urea utilization genes by Verrucomicrobia that we observed suggests that urea can be beneficial for bacterial species living under the ice in temperate and boreal lakes as well, given the similar constraints of temperature, light and organic nutrient availability.

### Implications

There is a growing acknowledgment among aquatic ecologists as to the need to study the full annual cycle of lakes in order to understand lake dynamics, particularly as lake temperature increases worldwide (O’Reilly *et al.*, 2015). This is exemplified by a recent quantitative synthesis on under ice ecology in which winter-summer patterns of nutrients, phytoplankton, and zooplankton were investigated (Hampton *et al.*, 2017) Though full-year time-series of these variables are rare, studies that also include microbial community structure and function variables are even more so although impressive long-term microbial time-series of the ice-free period do exist (Bendall *et al.*, 2016; Linz *et al.*, 2017). In addition to a few other studies, our findings substantiate the idea that understanding the under-ice microbiome is important for predicting the dynamics of seasonally ice-covered lakes in the future (Bertilsson *et al.*, 2013). Although this study was illuminating regarding under ice microbial metabolism, we focused solely on winter-associated Verrucomicrobia. Community wide winter-summer patterns remain to be elucidated and will likely vary tremendously between lakes. Hence, a much wider range of studies must be executed before generalizable patterns can be reported. Moreover, meta-omics studies such as this are, in essence, hypothesis-generating and future work that includes targeted enrichment/cultivation and *in situ* rate measurement-based approaches are required to validate and quantify microbial metabolic contributions to nutrient cycling in lake environments during winter.

## Methods

### Sampling

Microbial samples were collected from three freshwater lakes: Lake Croche (45°59’N, 74°01’W), Lake Montjoie (45°24’N, 72°14’W) and Lake Simoncouche (48°14’N, 71°15’W) in conjunction with the Groupe de Recherche Interuniversitaire en Limnologie et en Environnement Aquatique (GRIL) Monitoring Program. During 3 years (2013-2015), epilimnion and metalimnion samples were collected biweekly during ice-free periods and monthly during ice-covered periods of the year (see **Table S1** for a full description of samples collected for microbial analyses). In winter, samples were collected from just below the ice, as well as deeper in the water column for lakes Simoncouche and Croche, while a single integrated sample of water collected from multiple depths was collected from Montjoie. Lake water, collected in acid-washed bottles, was pre-filtered through 53 μm mesh, followed by sequential filtration on to a 3 μm polycarbonate filter to collect particle-attached cells followed by a 0.22 μm Sterivex filter to collect free-living cells. 1.8 ml of sucrose-based lysis buffer was added to samples collected for DNA extraction, 1.8 ml of RNA later was added to samples collected for RNA extraction, and filters were stored at −80° C until processing.

Water column profiles of environmental variables (**Table S2**) were measured including temperature, dissolved oxygen, pH, specific conductivity and oxidation potential, directly in the field using a multiparameter sonde (YSI, OH, USA). Water samples were collected at the same depths as the microbial samples. Analyses of total phosphorus (TP), total nitrogen (TN), total dissolved phosphorus (TDP), total dissolved nitrogen (TDN), dissolved organic carbon (DOC), nitrate (NO_3_^−^), nitrite (NO_2_^−^), ammonium (NH_4_^+^), ions (Cl^−^, PO_4_^3−^, SO_4_^2−^, Ca^2+^, Mg^2+^, Na^+^), and Chlorophyll *a* were performed in the GRIL laboratory at Université de Montréal (Montreal, Canada).

### Nucleic acid extraction

DNA was extracted from 0.22 μm Sterivex filters using a phenol/chloroform-method modified from Zhou (1996). Sterivex filters were thawed on ice and the storage buffer was removed. The storage buffer was concentrated into Amicon 30 kD filter (500 Hl at a time) followed by centrifugation for 20 minutes at 10,000 g. 500 μl storage buffer was repeatedly added until concentrated to a final volume of 100 μl. Buffer exchange was conducted twice by washing storage buffer with 500 μl of TENP buffer (600 mg Tris, 740 mg EDTA, 580 mg NaCl, 2 g Polyvinylpyrrolidon and 100 ml milliQ, pH 8). We then broke open the Sterivex filter and removed the filter. The filter was split into halves and placed inside a 2 ml Eppendorf tube. To conduct the cell lysis and digestion, 0.37 grams of 0.7 mm pre-sterilized Zirconium beads, 60 μl of 20 % SDS, 100 μl concentrated buffer exchanged filtrate, 500 μl TENP buffer and 500 μl phenol-chloroform-isoamylalcohol (PCI) 25:24:1 were added to the 2 ml Eppendorf tube containing the shredded filter. Then, the sample was vortexed for 10 minutes. Samples were incubated for 10 min in a 60 °C water bath followed by incubation on ice for 1 min. The samples were centrifuged for 6 min at 10,000 rpm and 4°C. Supernatant was transferred to a clean 1.5 ml Eppendorf tube, 500 μl phenol-chloroform-isoamylalcohol (PCI) 25:24:1 wasadded and samples were vortexed briefly. Then, samples were centrifuged for 6 min at 10,000 rpm and 4 °C. The supernatant was transferred to a new 1.5 ml Eppendorf tube. The PCR step was repeated until there was no longer any white precipitate at the interface (usually 2 times). DNA was precipitated by adding 120 μl of 3 M sodium acetate followed by 1 ml of 96% ethanol. DNA was precipitated at −20 °C for at least 1.5 hours, followed by centrifugation for 60 minutes at 13,000 rpm and 4 °C. The supernatant was decanted, and the pellet was washed with 850 μl of 80 % ethanol. We incubated samples for 10 minutes on ice followed by short vortexing and then centrifuging samples for 15 minutes at 13,000 rpm and 4 °C. The supernatant was removed and the DNA was resuspended in 50 μl TE or Tris-HCl (pH 7.5-8).

RNA was extracted from 0.22 μm Sterivex filters with a modified protocol (Shi *et al.*, 2009; Stewart *et al.*, 2010) which employs both the mirVana miRNA isolation kit (Invitrogen) and the RNeasy RNA cleanup kit (Qiagen). Samples were thawed and had the RNAlater (Invitrogen) surrounding the Sterivex filter removed (approximately 1700 μl) and discarded. 1700 μl of mirVana lysis buffer was added to the Sterivex filter and vortexed to lyse bacterial cells attached to the filter. Total RNA was then extracted from the lysate according to the mirVana protocol. Purified sample (100 ul) was treated with 2 ul DNase (New England Biotech) incubated at 65°C for 1-2 hours to remove genomic DNA, and concentrated using the RNeasy RNA cleanup kit (Qiagen).

### 16S rRNA gene sequencing and analysis

16S rRNA gene sequence data was generated from 143 samples (**Table S1**). The V3 region of the 16S rRNA gene was amplified using the universal primers (341F: 5’-CCTACGGGRSGC AGC AG-3’ and 515R: 5’-TTACCGCGGCKGCTGVC AC-3’) (Klindworth *et al.*, 2013). Two-step PCR reactions (modified from Berry et al., 2011) were conducted in 25 μl volume contained 0.5 μM MgCl_2_, 0.2 mM deoxynucleotide, 0.2 μM each primer and 1U of Phire Hot Start II DNA Polymerase (Finnzymes Thermo Fisher Scientific). The template was amplified using non-barcoded PCR primers for 20 cycles, followed by 1:50 dilution of the PCR product and 10 additional cycles of amplifications with barcoded PCR primers. The thermal program consisted of an initial 95 °C denaturation step for 4 min, a cycling program of 95 °C for 30 s, 52 °C for 30 s, and 72 °C for 60 s, and a final elongation step at 72 °C for 7 min. Reverse primers were barcoded with specific IonXpress sequence to identify samples. PCR products were purified using QIAquick Gel Extraction Kit (Qiagen), quantified using Quantifluor dsDNA System (Promega), pooled at equimolar concentration and sequenced using an Ion Torrent PGM system on a 316 chip with the ION Sequencing 200 kit as described in Sanschagrin and Yergeau (2014).

V3 region 16S rRNA sequences were analyzed using MOTHUR (Schloss *et al.*, 2009). Sequences with an average quality of <17, length <100 bp or that did not match the IonXpress barcode and both the PCR forward and reverse primer sequences were discarded. Potential chimeric sequences were identified using UCHIME (Edgar *et al.*, 2011) and discarded. Verrucomicrobia sequences were identified by taxonomic analysis in MOTHUR.

### Metagenome sequencing, assembly, annotation and binning

DNA sequencing of 24 samples (**Table S1**) was performed at the Department of Energy Joint Genome Institute (JGI) (Walnut Creek, CA, USA) on the HiSeq 2500-1TB (Illumina) platform. Paired-end sequences of 2 × 150bp were generated for all libraries. A combined assembly of all raw reads was generated using MEGAHIT (https://github.com/JGI-Bioinformatics/megahit) with kmer sizes of 23,43,63,83,103,123. Gene prediction and annotation was performed using the DoE JGI IMG/M functional annotation pipeline (Markowitz *et al.*, 2014). Metagenomic binning based on tetranucleotide frequency and differential coverage was performed with MetaWatt version 3.5.2 (Strous *et al.*, 2012) using the following settings: base pair cutoff of >2000 on the co-assembly, and a relative weight of binning coverage of 0.75. The identity of bins was assessed using the phylogenetic analysis of a concatenation of single copy core genes implemented in MetaWatt, MAFFT aligner and FastTreeMP for tree inference (Price *et al.*, 2010). Genome completeness and contamination was assessed with CheckM (Parks *et al.*, 2015), which relies on pplacer (Matsen *et al*, 2010), prodigal (Hyatt *et al.*, 2012) and HMM (Eddy, 2011). Fourteen MAGS with low contamination (<5%) and substantially or near-complete (>70%) were obtained, as described by the bin quality terminology proposed in Parks *et al.* 2015. Despite a completeness value of 58%, we also included Opi-242, given its abundance during the ice-covered period of Lake Simoncouche.

### Ecological association of Verrucomicrobia bins

We performed a canonical constrained analysis (CCA) using the *vegan* package (Oksanen *et al.*, 2017) in R (R Core Development Team, 2016). The 20 environmental variables measured were tested for normality using a Shapiro test, and transformed using *powerTransform*, followed by a Box-Cox transformation *(bcPower)*, using the *car* package (Fox *et al.*, 2016) for all variables for which p<0.05 in the Shapiro test. A linear model was fitted to each pair of variable, and we removed variables for which the correlation coefficient was greater than 0.70 from the CCA. In brief, we performed the CCA using 14 of the environmental variables, and the complete species matrix containing 24 samples and 54 MAG.

### Phylogeny of Verrucomicrobia

A concatenated gene phylogeny tree was created based on 4 of the 5 genes as in He *et al.* (2017), in which the 15 Quebec MAGs were put in the contact of the 19 MAGs from Lake Mendota and Trout Bog (Wisconsin), and 7 MAGs from ice-covered Lake Baikal (Cabello-Yeves, Zemskaya, *et al.*, 2017) and 17 MAGs from the Tous and Amadorio reservoirs (Cabello-Yeves, Ghai, *et al.*, 2017). Because none of the 15 MAGs in our study contained the DNA polymerase III beta subunit (but had other subunits such as alpha), the concatenated gene phylogeny contained instead the following four genes: TIGR01391 (DNA primase), TIGR01011 (Small subunit ribosomal protein S2), TIGR00460 (Methionyl-tRNA-formyltransferase), and TIGR00362 (Chromosomal replication initiation factor). 17 out of the 18 MAGs from the Tous and Amadorio Reservoirs had these genes. To create the phylogenetic tree, sequences for each gene were aligned using MUSCLE (Edgar, 2004) and a maximum-likelihood tree was created for each individual gene to ensure that these proteins were conserved and represented the phylogenetic relationships between groups, before concatenation of the sequences using Mesquite (Maddison & Maddison, 2017). The confidence score of each amino acid position in the multiple sequence alignment was calculated using ZORRO (Wu *et al.*, 2012), and amino acid position that had a score below 0.5 was manually deleted using Mesquite (Maddison & Maddison, 2017) to obtain a more accurate phylogenetic inference. MEGA6.06 (Tamura *et al.*, 2013) was used to generate a maximum likelihood phylogeny of 65 taxa, using a bootstrap of 100 iterations. The substitution model was the Jones-Taylor-Thornton (JTT) model for amino acids. The rates among site were Gamma distribution, with 4 gamma categories. The maximum-likelihood (ML) heuristic method was Nearest-Neighbour Interchange (NNI), and the initial tree was NJ. Finally, the branch swap filter was Very Strong. Despite that four instead of five genes were used, the phylogenetic structure of the tree (and relationships between taxa) is the same as when using 5 genes, as in He *et al.* (2017). In order to include all 18 MAGs from (Cabello-Yeves, Ghai, *et al.*, 2017), an automated concatenated gene phylogeny was created using PhyloPhlan (Segata *et al.*, 2013) (**Figure S2**). The phylogenetic structure of the PhyloPhlan tree was consistent with the manually curated tree (**Figure 3**).

### Comparative genomics and functional annotation

The distribution of protein-encoding gene content between genomes was determined using proteinortho (Lechner *et al.*, 2011). Inference of protein function and metabolic reconstruction was based on the IMG annotations provided by the JGI, including KEGG, Pfam, EC numbers, and Metacyc annotations. Metabolic reconstruction was also facilitated by generated pathway genome databases for each MAG using the pathologic software available through Pathway Tools (Karp *et al.*, 2009). In addition, we annotated carbohydrate-active enzymes using dbCan (Yin *et al.*, 2012) and hydrogenase classes using HydDB (Sondergaard *et al.*, 2016).

### Metatranscriptomic analysis of Verrucomicrobia gene expression patterns

cDNA library preparation and sequencing of 24 samples (**Table S1**) was performed at the Department of Energy Joint Genome Institute (JGI) (Walnut Creek, CA, USA) on the HiSeq 2500-1TB (Illumina) platform. Paired-end sequences of 2 × 150 bp were generated for all libraries. The metatranscriptome dataset comprises of 24 samples (6 Croche, 7 Montjoie and 11 Simoncouche). BBMAP (https://jgi.doe.gov/data-and-tools/bbtools/bb-tools-user-guide/bbmap-guide/) was used to map the raw quality filtered reads to the CDS in each 15 Verrucomicrobia MAG, using the option *min_id*=0.97 instead of the default one of 0.7. The RPKB, recruitments per kilo bases of billion reads per sample, was calculated instead of the RPKM (recruitments per kilo bases of million mapped reads) to control for differences in raw reads between samples.

### ANI and AAI calculations and fragment recruitment

ANI values were calculated with ANICalculator (https://ani.jgi-psf.org/html/anicalculator.php) using the default settings, but the alignment fraction was too low (median 0.02-0.03 %) to produce meaningful results. Therefore, we assessed genome similarity using AAI values calculated using CompareM (https://github.com/dparks1134/CompareM) and the default settings. Fragment recruitment was performed using BBMAP (https://jgi.doe.gov/data-and-tools/bbtools/bb-tools-user-guide/bbmap-guide/) and the following settings: *minid*=94, *maxlen*=500, *idtag*=t. To create the heatmap of relative coverage values, we divided the number of reads mapped by the size of the metagenome for each MAG.

### Data accessibility

The 24 metagenomes and the associated co-assembly can be downloaded at the JGI (IMG Genome ID: 3300010885). The 24 metatranscriptome and the associated coassembly can be downloaded at IMG Genome ID: 3300013295. MAG data is available at NCBI under the accession numbers XXX-XXX

## Acknowledgments

We would like to thank Morgan Botrel and Stéphanie Massé for coordinating the Lake Sentinels Project, a program funded by the Groupe de recherche interuniversitaire en limnologie et en environnement aquatique (GRIL), and for sampling the Lakes Croche and Montjoie. We thank Guillaume Grosbois, Tobias Schneider and Maxime Wauthy for sampling the Lake Simoncouche. Funding from the Canadian Natural Sciences and Engineering Research Council (NSERC) Discovery Grants (D.W., M.R., B.B., and Y.H.) and the Canada Research Chair Program (D.W., M.R., Y.H.) are also acknowledged. The work conducted by the U.S. Department of Energy Joint Genome Institute, a DOE Office of Science User Facility, is supported under Contract No. DE-AC02-05CH11231. P.T. was funded by NSERC-CREATE ÉcoLac, and Fonds de Recherche du Québec Nature et Technologie (FRQNT).

## Supplementary Figures and Tables Titles

**Figure S1.**
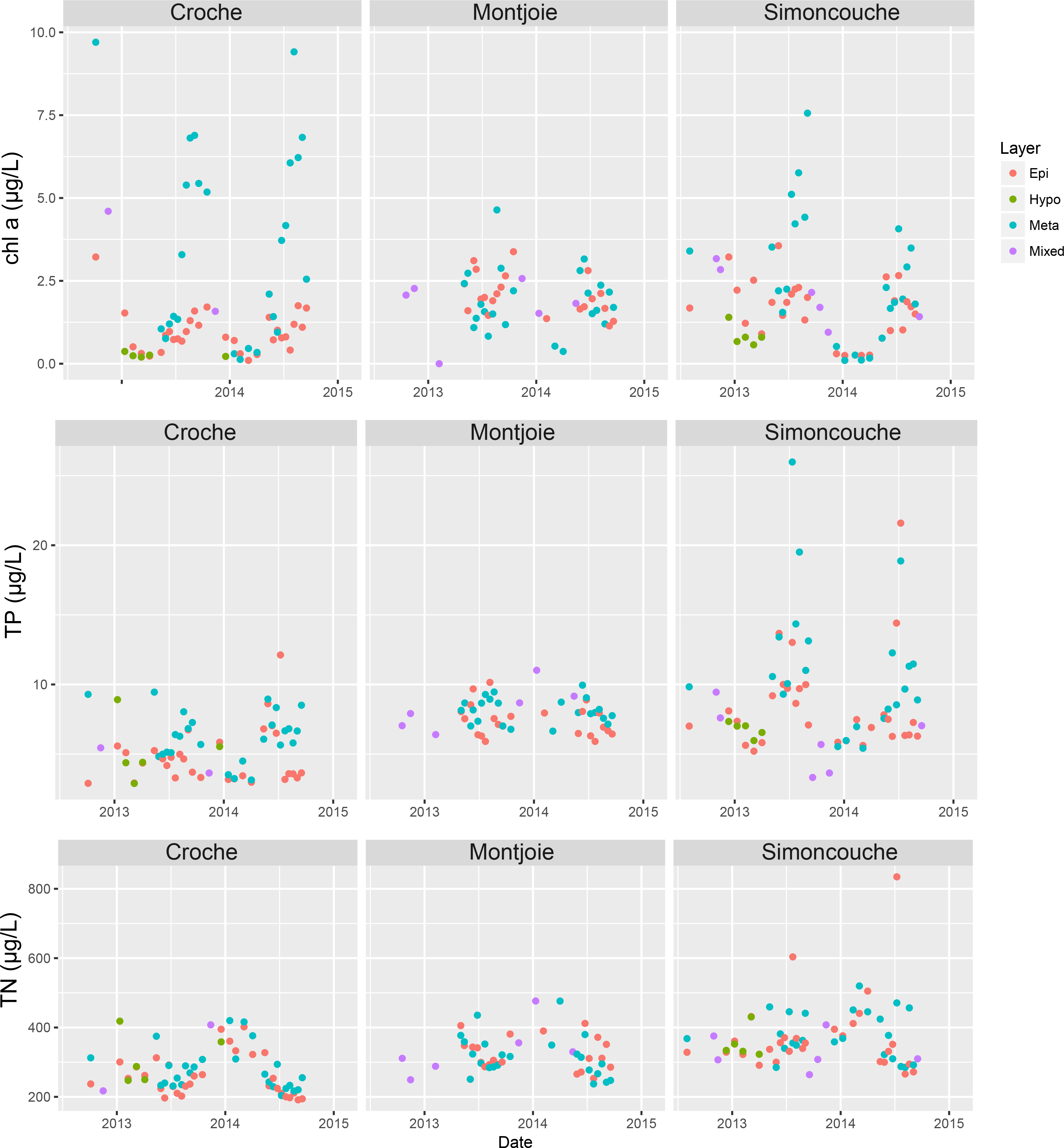
Scatter plots of chlorophyll *a*, total phosphorus (TP) and total nitrogen (TN) coloured by strata over the period of the study in the three Quebec lakes.

**Figure S2.**
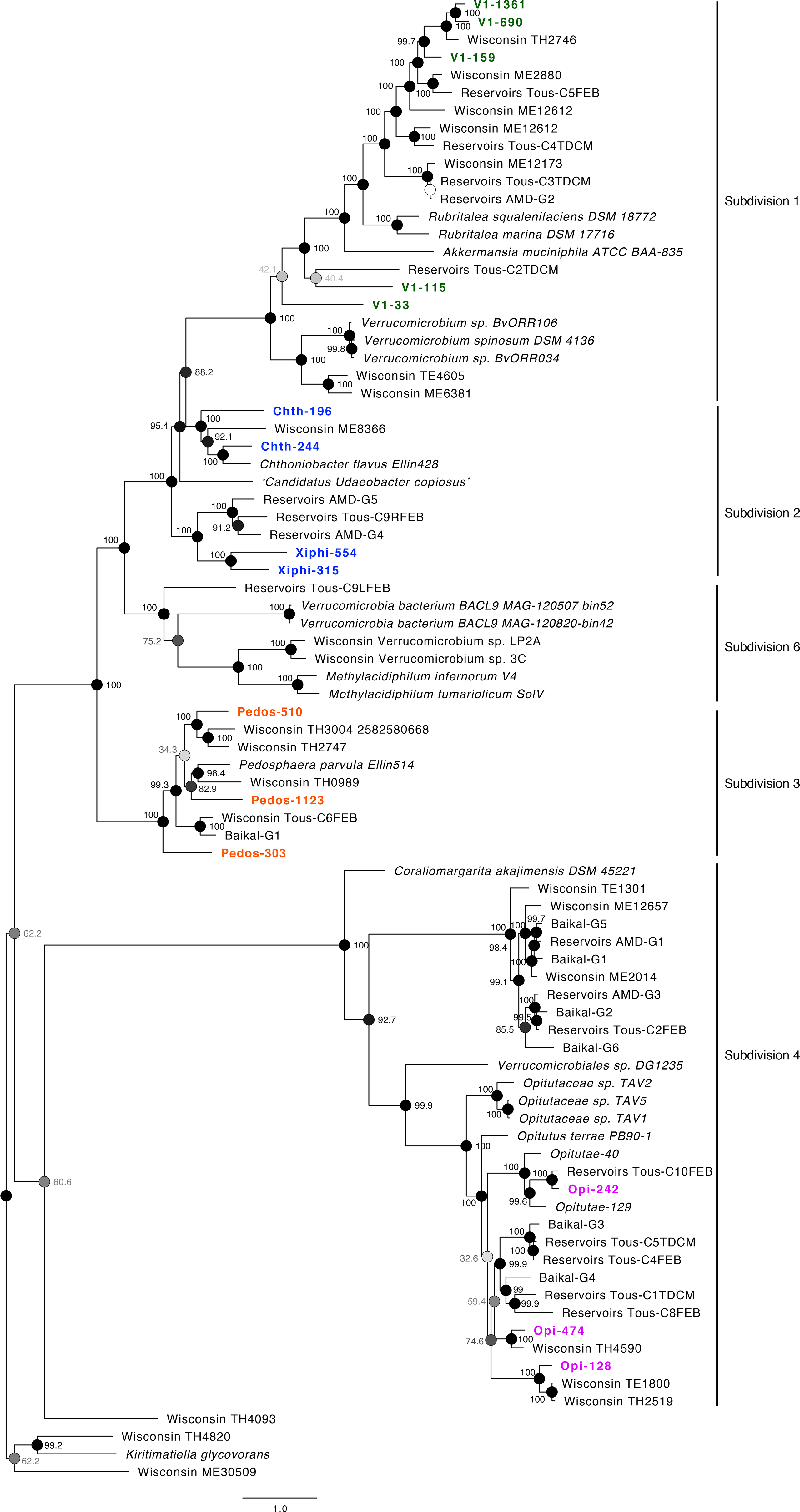
Concatenated gene phylogeny using Phylophlan showing the 15 Quebec MAGs with all currently existing Verrucomicrobia MAGs from Lake Baikal, Trout Bog, Lake Mendota, Tous and Amadorio reservoirs.

**Table S1.** Description of the 16S rRNA gene, metagenome and metatranscriptome datasets employed in this study.

**Table S2.** Environmental variables measured in the field and in the laboratory. Abbreviations are the same as those used in Figures 1B and 2B.

**Table S3.** Pair-wise amino acid identities (AAI) comparisons between all available freshwater Verrucomicrobia MAGs, divided into subdivisions.

**Table S4.** Fragment recruitment results of 15 MAGs against 22 freshwater metagenomes.

**Table S5.** Expression values (RPKB) for GHs in each of the 15 MAGs.

